# Classification of Extracellular Vesicles based on Surface Glycan Structures by Spongy-like Separation Media

**DOI:** 10.1101/2022.05.10.491426

**Authors:** Eisuke Kanao, Shuntaro Wada, Hiroshi Nishida, Takuya Kubo, Tetsuya Tanigawa, Koshi Imami, Asako Shimoda, Kaori Umezaki, Yoshihiro Sasaki, Kazunari Akiyoshi, Jun Adachi, Koji Otsuka, Yasushi Ishihama

**Affiliations:** Graduate School of Pharmaceutical Sciences, Kyoto University, Sakyo-ku, Kyoto 606-8501, Japan; National Institutes of Biomedical Innovation, Health and Nutrition, Ibaraki, Osaka 567-0085, Japan; Department of Material Chemistry, Graduate School of Engineering, Kyoto University, Katsura, Nishikyo-ku, Kyoto 615-8510, Japan; Precursory Research for Embryonic Science and Technology (PRESTO), Japan Science and Technology Agency (JST), 4-1-8 Honcho, Kawaguchi, Saitama 332-0012, Japan; Department of Polymer Chemistry, Graduate School of Engineering, Kyoto University, Katsura, Nishikyo-ku, Kyoto 615-8510, Japan

**Keywords:** extracellular vesicles, lectin affinity chromatography, heterogenity, surface glycan structure, proteome analysis

## Abstract

Extracellular vesicles (EVs) are lipid bilayer vesicles that enclose various biomolecules. EVs hold promise as sensitive biomarkers to detect and monitor various diseases. However, they have heterogenous molecular compositions. The compositions of EVs from identical donor cells obtained using the same purification methods may differ, which is a significant obstacle for elucidating objective biological functions. Herein the potential of a novel lectin-based affinity chromatography (LAC) method to classify EVs based on their glycan structures is demonstrated. The proposed method utilizes a spongy-like monolithic polymer (spongy monolith, SPM), which consists of poly(ethylene-co-glycidyl methacrylate) with continuous micropores and allows an efficient *in-situ* protein reaction with epoxy groups. Two distinct lectins with different specificities, *Sambucus sieboldiana agglutinin* and concanavalin A, are effectively immobilized on SPM without impacting the binding activity. Moreover, high recovery rates of liposomal nanoparticles as a model of EVs are achieved due to the large flow-through pores (>10 μm) of SPM. Finally, lectin-immobilized SPMs are employed to classify EVs based on the surface glycan structures and demonstrate different subpopulations by proteome profiling.

## 1. Introduction

Extracellular vesicles (EVs) are lipid spheres formed from the plasma membranes and are released by living cells. EVs encase endogenous bio-related molecules such as *mi*RNA, proteins, lipids, and glycans.^[1–3]^ Small EVs (30–150 nm), called exosomes, act as an essential mediator of intercellular communications. They help regulate several key physiological processes that keep our bodies healthy by delivering cargo into the cytoplasm of the recipient cells.^[4]^ EVs have potential as biomarkers for cancers and neuropathic diseases because these components reflect the donor cell states.^[5–8]^ Simultaneously, they hold promise as biological sources of drug delivery systems for therapeutics or vaccine production systems in the field of drug discovery.^[9–11]^

Despite their potential in clinical applications, the heterogeneity of the EV molecular composition is a risk against authenticity for studies on functions.^[12, 13]^ For example, although various tetraspanins with a wide cellular expression (CD9, CD63, CD81, and CD82) are general marker proteins on EV membranes, the tetraspanin content of the EVs produced from identical cells may differ.^[14, 15]^ Kugeratski et al. reported that EVs collected by density gradient (DG) size-exclusion chromatography (SEC) or ultracentrifugation (UC) show a heterogeneous abundance of tetraspanins and highlighted the potential of syntenin-1 as a putative universal marker of EVs.^[16]^ Another example demonstrated that although heterogeneous, phosphatidylserine (PS) can be used as an EV marker for immunoaffinity methods. However, PS-enriched EVs are characterized by a lower density, a larger size, and a more negative zeta potential than those collected by DG.^[17, 18]^ These membrane components play essential roles in targeting. Several *in vivo* studies have investigated the uptake of EVs.^[19–25]^ Therefore, classification methods of EVs based on the membrane components are urgently required to standardize their precise functions for clinical applications.

Membrane glycoproteins and glycolipids cover the surface layers of cells.^[26, 27]^ Their glycan patterns depend on the cell type and the condition. These patterns are associated with biological and pathological processes. Glycans on the surfaces of EVs should affect cellular interactions, and biodistribution interrogation of the glycosylation characteristics of EVs is needed.^[28–32]^ Lectins, which are non-antibody carbohydrate-binding proteins, recognize particular glycans commonly found on the surfaces of cells and vesicles. Lectins have been utilized for detailed profiling of EV glycosylations.^[33]^ Previously, our study utilizing evanescent field fluorescence-assisted lectin microarray reported that the glycan pattern of EVs depends on the osteogenic stages.^[34]^ Furthermore, glycoengineering showed specific uptake behavior into cells for EVs with remodeled surface glycan patterns.^[35, 36]^These reports suggest that glycans have potential as novel standardized indicators of the heterogeneity of EVs.

Lectin affinity chromatography (LAC) has been utilized in purification procedures based on the glycan structure in the biomedical field.^[37–40]^ Compared to bead-based adsorption, chromatographic separation is faster and more efficient with a higher reproducibility. However, few studies have evaluated LAC separation of liposomal nanoparticles larger than 100 nm, which includes EVs. Although agarose and silica particles are valuable materials for lectin immobilization, their narrow interparticle volumes may promote clogging of nanoparticles.^[41–43]^

Our group previously reported a spongy-like monolithic separation media (spongy monolith, SPM).^[44]^ SPM consists of poly(ethylene-co-glycidyl methacrylate) (PEGM) and displays a superior permeability, operability, and cost performance. Due to the continuous three-dimensional structures with large flow-through pores (>10-μm diameter), SPM realizes an efficient *in-situ* reaction between the epoxy groups. Hence, SPM may be suitable as an LAC media of liposomal nanoparticles, including EVs.

In this study, we develop a novel classification method for EVs using lectin-immobilized SPMs and demonstrate the heterogeneity of EVs based on the difference in surface glycans. Two distinct lectins with different specificities, concanavalin A (ConA) and *Sambucus sieboldiana agglutinin* (SSA), are immobilized onto the SPM media *in-situ*, and their affinity reactions are quantitatively examined with glycoproteins and mannose-labeled liposome as a model vesicle for EVs. Finally, the lectin-immobilized SPMs are used to classify small EVs based on their surface glycan structures and the differences in the proteome profiles of the collected EVs are analyzed by nano-flow liquid chromatography-tandem mass spectrometry (LC/MS/MS).

## 2. Results and discussion

### 2.1. Separation behaviors of glycoproteins and mannose-labeled liposome on lectin-immobilized SPMs

Two different lectins, ConA specific to high-mannose glycans and SSA specific to *α*-2,6-sialylated glycans, were immobilized on SPMs as models of LAC columns (ConA-SPM and SSA-SPM). Glucose oxidase (GOx) with high-mannose glycans as well as transferrin (Tf) with α-2,6-sialylated glycans were empoyed to evaluate the separation behavior. Figure 1 shows the chromatogram on each column. Glycoproteins were effectively retained on their respective lectin-immobilized SPMs and were eluted with extremely high concentrations of hapten sugars. Hence, lectins were successfully immobilized on SPMs while retaining their specific binding activities for their respective glycans.

**Figure 1.**
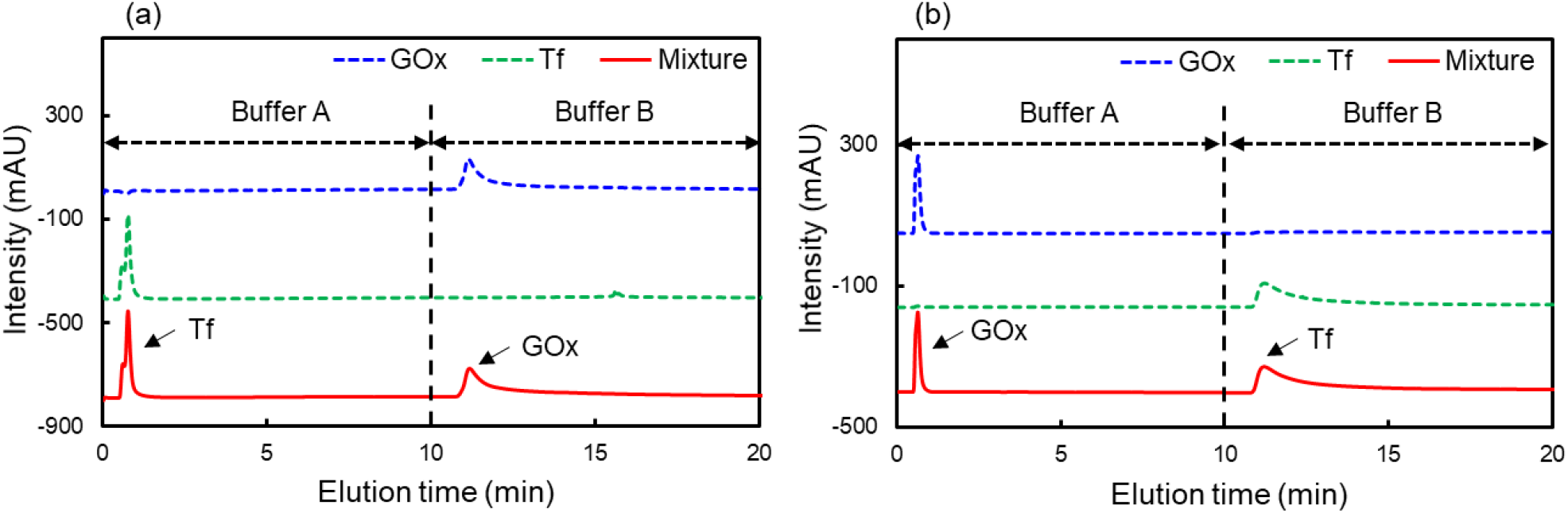
Lectin affinity chromatography of glycoproteins on (a) ConA-SPM and (b) SSA-SPM. LC conditions: column, (a) ConA-SPM (50 mm × 4.6 mm I.D.), (b) SSA-SPM (50 mm × 4.6 mm I.D.); detection, UV 280 nm; flow rate, 0.5 mL/min; mobile phase, buffer A (20 mM HEPES buffer (pH 7.4) with 150 mM NaCl, 1 mM MgCl_2_, 1 mM CaCl_2_, and 1 mM MnCl_2_), and buffer B (200 mM hapten sugar in buffer A).

Next, the loading capacities for the respective glycoproteins were investigated (Fig. S1). Both columns showed tailing peaks derived from overloading glycoproteins in the flow-through fractions (Figs. S1a,b, Supporting Information). Increasing the sample loading led to larger peak areas of the flow-through and hapten-elution fractions (Figs. S1c,d, Supporting Information). Furthermore, the total peak area of both fractions and the loading sample amount had linear relationships with *R*^2^ > 0.99. Thus, the LAC separation was achieved without nonspecific adsorption derived from high concentration samples.^[45]^ The estimated loading capacities of ConA-SPM and SSA-SPM toward the glycoproteins were 3.53 nmol and ×.×× nmol, respectively.

The designed LAC method was directly utilized to separate mannose-labeled liposomes (Fig. S2a, Supporting Information). The liposomes were labeled with rhodamine for easy detection by fluorescence spectrometry in the LC system. The average particle size, which was measured by dynamic light scattering (DLS), was ca. 137.3 nm (Fig. S2b, Supporting Information). A colorimetric lipid quantitation kit indicated that the final 1,2-dioleoyl-sn-glycero-3-phosphocholine (DOPC) concentration in the liposome solution was 217.4 mM. Since DOPC and PA-PEG-mannose were mixed at a 100:1 molar ratio, the liposome solution contained 1.24 mM mannose structures. The liposome solution (5 μL), containing 10.9 nmol of mannose structures, was injected into the ConA-SPM column to assess the adsorption capacity and recovery rate. For comparison, a commercially available ConA-immobilized agarose particle (Concanavalin A, immobilized on Sepharose®4B, Sigma Aldrich Japan) was packed into an empty column (ConA-agarose column) and tested under the same conditions. Figure 2a shows the chromatograms of mannose-labeled liposomes on each column. Both columns retained a portion of the injected liposomes due to the affinity of ConA with their mannose structures.

**Figure 2.**
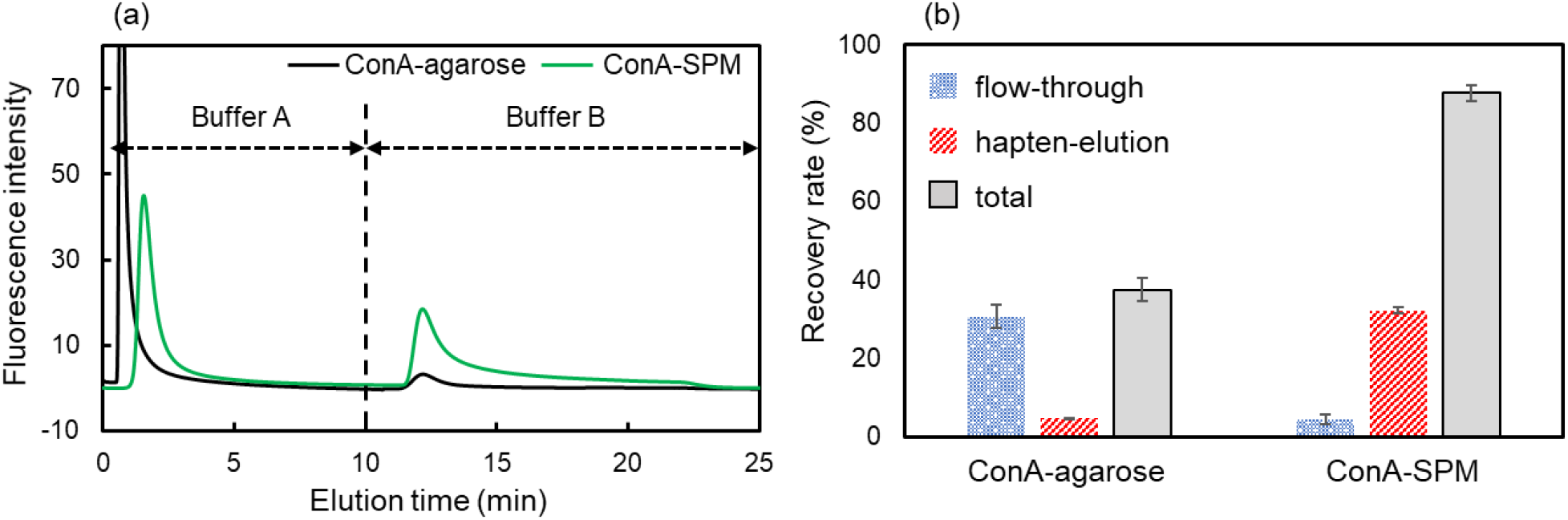
LAC of liposomal nanoparticles on the ConA-SPM and ConA-agarose columns. (a) Chromatograms of mannose-labeled liposomes, (b) recovery rate of the injected mannose-labeled liposomes in their respective fractions and the total on the ConA-SPM and ConA-agarose columns. Error bar amplitude matches the mean ± SD of three trials. LC conditions: column, ConA-SPM (50 mm × 4.6 mm I.D.), ConA-agarose (50 mm × 4.6 mm I.D.); detection, fluorescence E_x_, 560 nm, E_m_; flow rate, 0.5 mL/min; mobile phase, buffer A (20 mM HEPES buffer (pH7.4) with 150 mM NaCl, 1 mM MgCl_2_, 1 mM CaCl_2_, and 1 mM MnCl_2_), and buffer B (200 mM hapten sugar in buffer A).

Figure 2b summarizes the calculated recovery rates of the liposomes in both fractions and the total recovery rates. The calibration curve of the liposomes was prepared by the flow fluorescence detector in the LC system without an analytical column (Fig. S3, Supporting Information). Interestingly, the recovery rates of both flow-through and hapten-elution fractions were more significant on the ConA-SPM column than those on the ConA-agarose column. The number of mannose structures on the liposomes retained on ConA-SPM was roughly estimated as 3.3 nmol by the adsorption capacity toward GOx. In addition, the total recovery rates on the ConA-SPM column were 87.6%, and the injected liposomes were almost completely recovered, even though the total recovery rates were only 32.2% on the ConA-agarose column. Generally, packed materials must have sufficiently large pores for liposomal nanoparticles (more than five times) to interact with bio-ligands in stationary phases because they may become clogged and easily collapse due to their large size.^[46–48]^ Even though Sepharose®4B is a relatively large agarose matrix with bead sizes of 45–165 μm and a pore size of 42 nm,^[49]^ the particle size was not suited for non-destructive separation of the liposomes. In contrast, the average pore size of SPM, which was determined by a mercury porosimeter, was ~10 μm. Mesopores were not detected by nitrogen-gas adsorption analysis.^[44]^ Therefore, the mannose structures interacted efficiently with the ConA immobilized on the SPM surface, and liposomal nanoparticles could be separated nondestructively.

### 2.2. Suppression of nonspecific EV adoption to SPM by a blocking treatment

Issues in EV research include nonspecific adsorption of EVs to hydrophobic materials and pronounced particle losses. These are more likely to occur for purified vesicles.^[50–52]^ However, there is not a consensus on a method to suppress nonspecific adsorption of EVs. Here, we consider a blocking treatment with excess protein or hydrophilic polymers as a practical solution to prevent nonspecific binding in various bioanalytical methods.

Specifically, a protein A-immobilized SPM packed in an SPE cartridge was coated with various blocking agents. The recovery rates of EVs from SPM were evaluated by the fluorescence intensity (FI) of each fraction. Figure 3 shows blocking agents and recovery rates. The protein A-immobilized SPM was immersed in the blocking solution for 1 h and washed with a HEPES buffer (5 mL). EV samples (1 mL, protein content in 10.2 μg), which were collected by ultracentrifugation from the culture supernatant of HEK293sus, were passed through the SPM (flow-through fraction) and washed with HEPES buffer (1 mL × 4, wash fraction 1~4). The recovery rates were calculated as

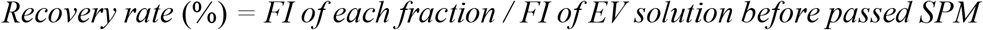

**Figure 3.**
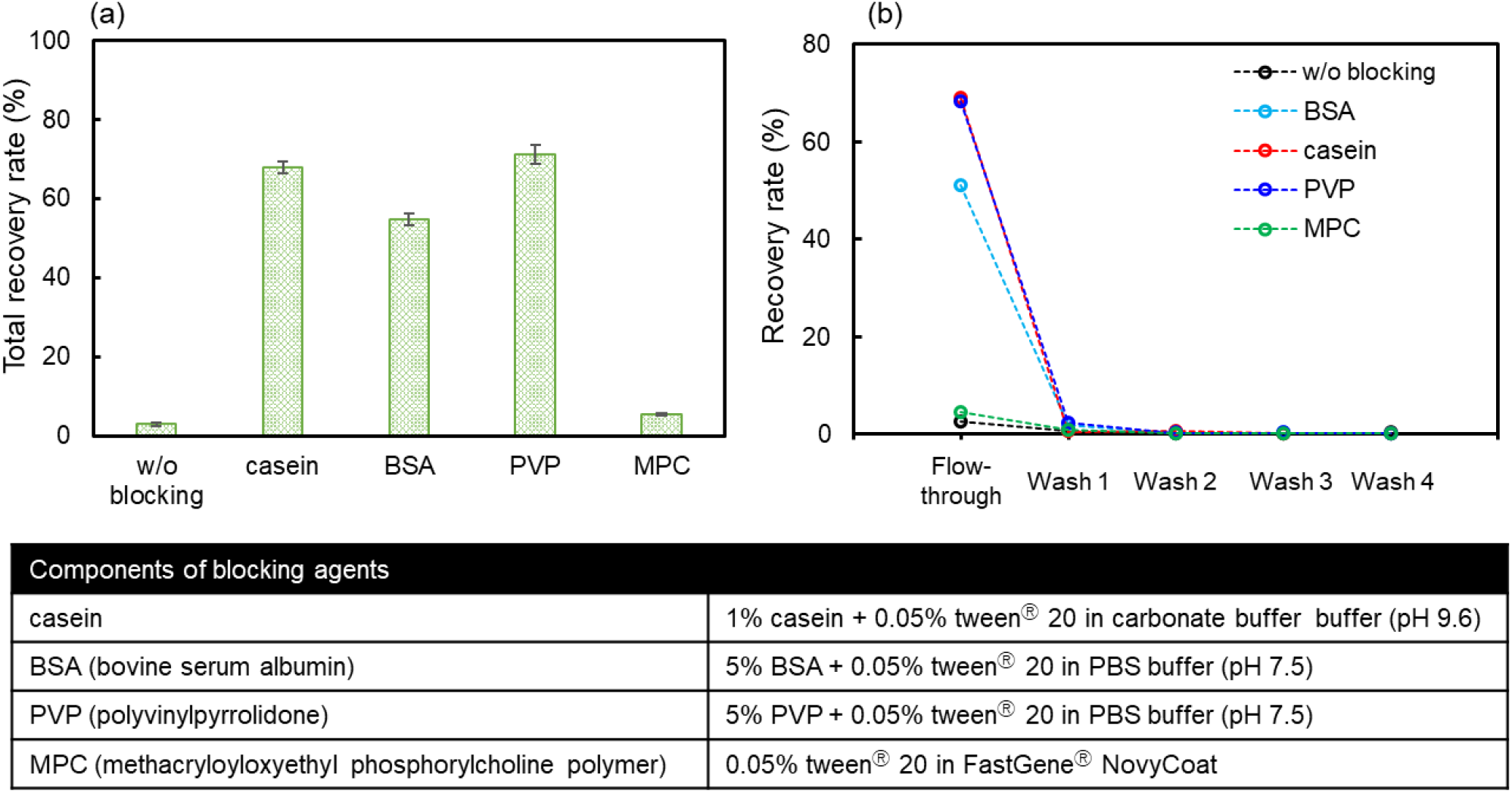
Recovery rates of EVs from protein A-immobilized SPM treated with various blocking agents. (a) Total recovery rates (error bar amplitude matches the mean± SD of three trials) and (b) recovery rates in each fraction. Table summarizes the blocking agent components.

Although the total recovery rates were less than 5% on SPM without blocking, casein and polyvinylpyrrolidone (PVP) effectively suppressed nonspecific adsorption on SPM (Fig. 3a, over 70%). The washing fraction contained tiny EVs, suggesting that they passed through without being retained on SPM (Fig. 3b). Surprisingly, a 2-methacryloyloxyethyl phosphorylcholine polymer solution (FastGene®NovyCoat, Nippon Genetics, Tokyo, Japan), which is a famous biomembrane-like polymer that suppresses protein adsorption,^[53]^ did not work as an effective blocking agent for SPM.

We employed PVP as the blocking agent to the lectin-immobilized SPM. The proteins were not suitable for blocking agents as they interfered with the later proteomics analysis of EVs. PVP blocking did not compromise the specific affinities, and LAC separation of the respective glycoproteins was achieved on both the lectin-immobilized SPMs after the blocking treatment with PVP (Fig. S4, Supporting Information).

### 2.3. LAC separation of EVs and their proteome profiling

We verified the effectiveness of our SPMs as an LAC platform for liposomal nanoparticles. The PVP blocking treatment suppressed nonspecific EV adsorption on SPMs. Then we carried out LAC separation of EVs on lectin-immobilized SPMs and proteome profiling of the separated EVs to determine the heterogeneity of EVs based on their surface glycan structures. Small ECs were collected by UC because EVs have both size-dependent heterogeneity, which ranges from nanometers to micrometers, and different components. This approach provided a relatively homogenous size distribution. The resulting EVs were characterized by their size and the expression of EV marker proteins by nano tracking analysis and western blotting analysis (Fig. S5, Supporting Information), respectively. Finally, a narrow particle size distribution (147.3 ± 2.1 nm) and expression of EV marker proteins (CD63 and HSC70) were confirmed.

Figures 4a,b show representative chromatograms of EVs on each lectin-immobilized SPM with different injection amounts of EVs. The bicinchoninic acid (BCA) assay determined the injected EV amounts were 5 (flow-through), 10 (hapten), and 20 μg (total). The small peaks were probably derived from partially injected EVs. This result suggests that lectin affinity reactions of EVs occur on both lectin-immobilized SPMs. The peak areas in both fractions were calculated to investigate the adsorption capacity (Figs. 4c,d). Minimal peaks were observed in the flow-through and hapten-elution fractions with 5 μg EVs on the respective SPMs. The peak areas in hapten-elution fractions significantly increased between 5 and 10 μg of EVs, whereas the peak areas did not increase significantly between 10 and 20 μg. These results indicate a loading capacity between 10 and 20 μg was achieved due to the saturation of lectin-immobilized SPMs. Therefore, the estimated appropriate amount of EV to these SPMs was 10 μg of protein equivalent. In addition, the size distributions after collecting the flow-through and hapten-elution fractions were investigated by nano tracking analysis. After passing lectin-immobilized SPMs, the morphological integrity of the EVs did not change (Figs. 5a–d). The mean diameter was 120–140 nm with a homogenous size distribution. Thus, LAC separation of EVs was achieved nondestructively. Furthermore, the particle number in each fraction and the total recovery rates were also evaluated by nano tracking analysis (Fig. 5e). The hapten-elution fractions on ConA-SPM and SSA-SPM contained 9% and 12% of the injected EVs, respectively. Additionally, the total recovery rates were 66% (ConA-SPM) and 58% (SS-SPM).

**Figure 4.**
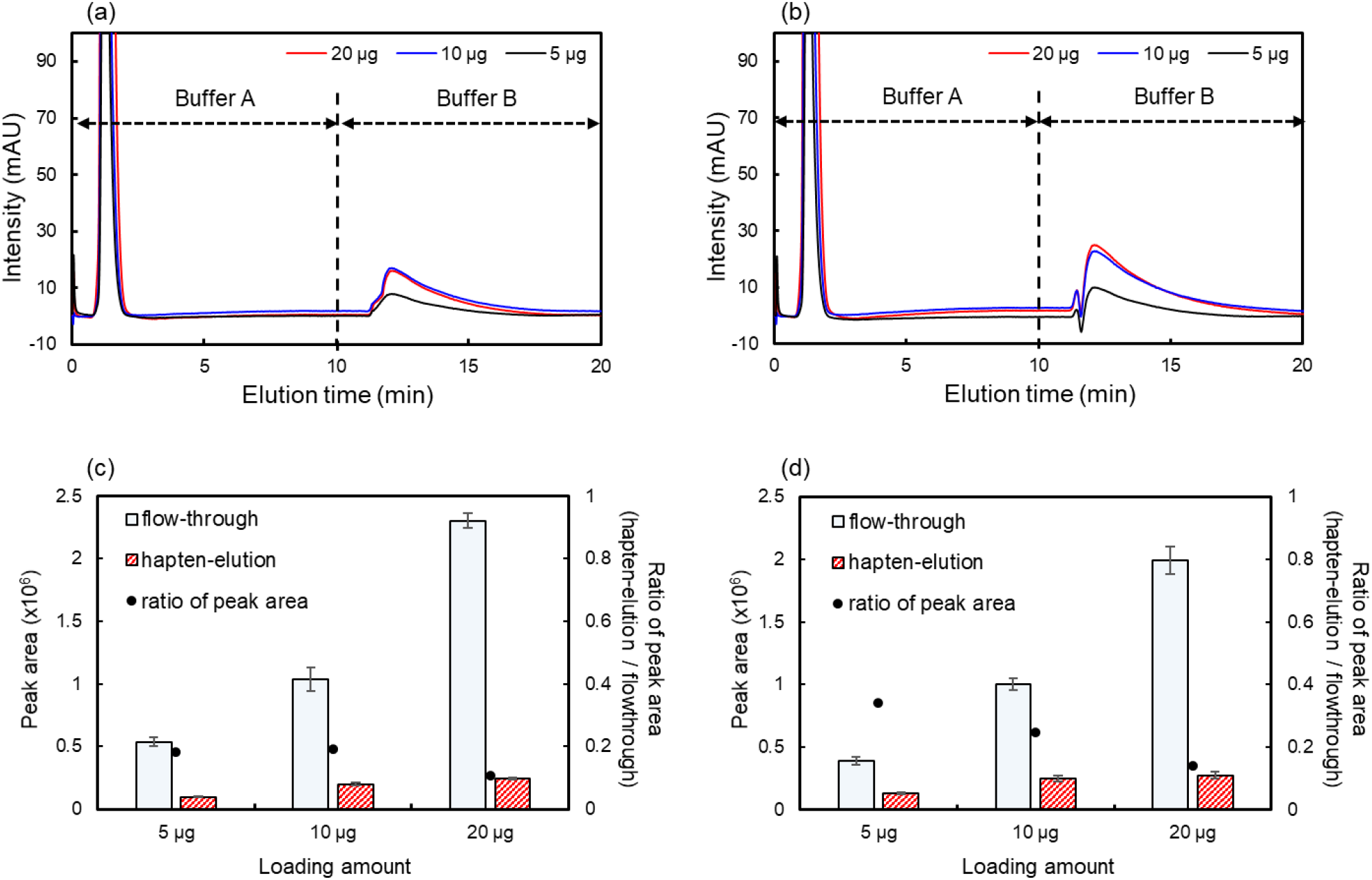
LAC separations of EVs collected by UC from HEK293sus on the lectin-immobilized SPMs. Chromatograms of EVs on (a) ConA-SPM and (b) SSA-SPM. Peak areas of flow-through and hapten-elution fractions, and their ratio on (c) ConA-SPM and (d) SSA-SPM (error bar amplitude matches the mean ± SD of three trials). LC conditions: column, (a), (c) ConA-SPM (50 mm × 4.6 mm I.D.), (b), (d) SSA-SPM (50 mm × 4.6 mm I.D.); detection, UV 280nm; buffer A (20 mM HEPES buffer (pH7.4) with 150 mM NaCl, 1 mM MgCl_2_, 1 mM CaCl_2_, and 1 mM MnCl_2_), buffer B (200 mM hapten sugar in buffer A)

**Figure 5.**
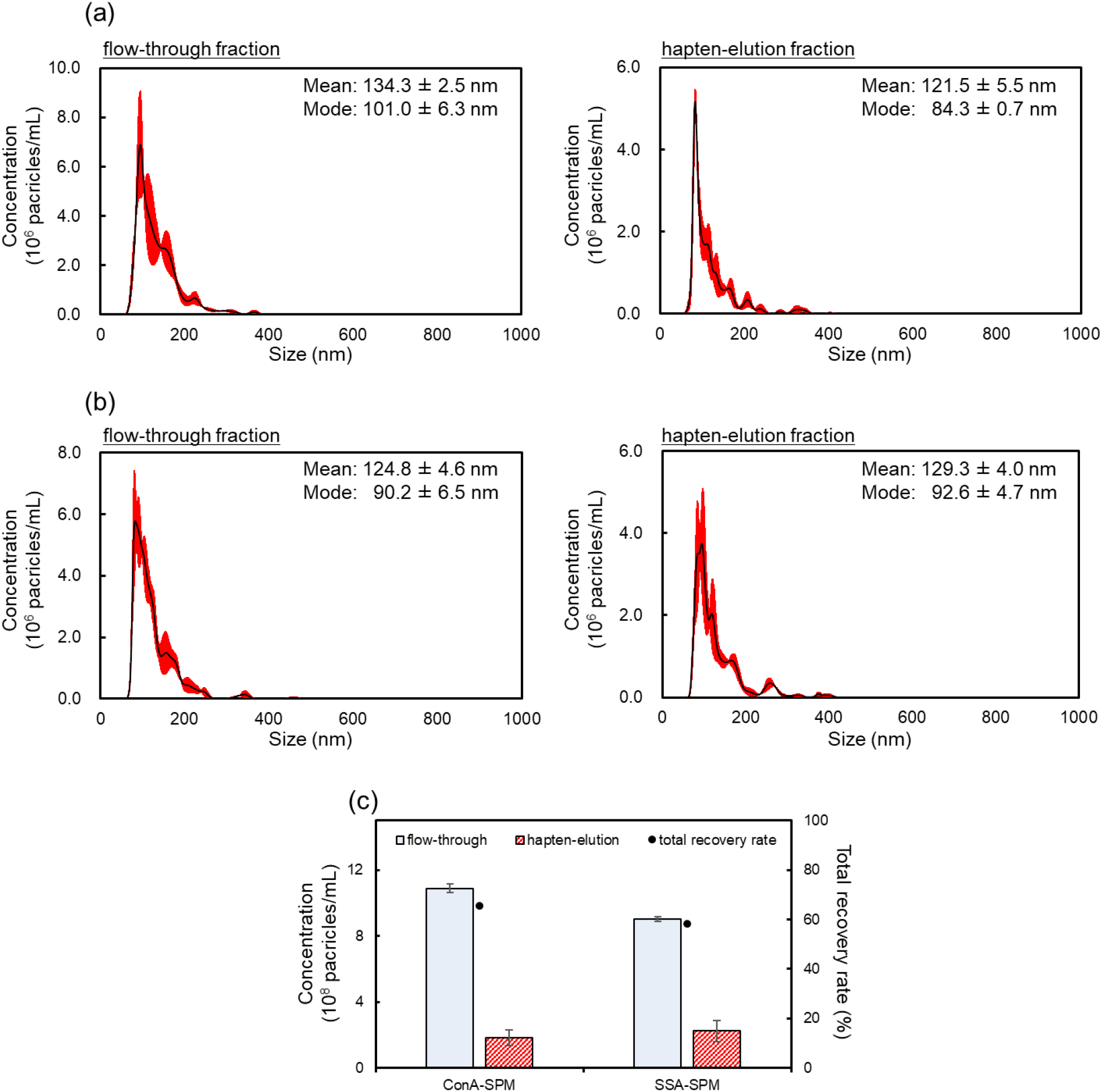
Size distribution eluted from (a) ConA-SPM and (b) SSA-SPM. (right) EVs in the flow-through fractions and (left) EVs in the hapten-elution fractions. (c) Particle concentration in each fraction, and the total recovery rate estimated from the injected EV particle number. Error bar amplitude matches the mean ± SD of three trials.

About half of EVs were lost during LAC analysis despite the blocking treatment with PVP. This loss may be due to the nonspecific adsorption on the LC system and EV storage tips along with lectin-immobilized SPMs.

Finally, the digested peptides from the EVs separated on SPMs were used for LC/MS/MS analysis to compare their proteome profiles. The LAC experiments were repeated three times with three aliquots. Then the injected amount of peptides for a single LC/MS/MS analysis was determined on NanoDrop.

The injected amounts were 0.82 μg and 0.76 μg peptides in flow-through fractions, and 0.39 μg and 0.42 μg peptides in hapten-elution fractions on ConA-SPM and SSA-SPM, respectively. For comparison, 1.04 μg peptides of the original EVs were also measured. Venn diagrams compared the lists of identified proteins from the original EVs and in each fraction (Figs 6a,b). Here, unique proteins identified in at least two of the three trials were used to increase the reliability. The commonly identified proteins in all samples were included in several EV markers such as tetraspanins (CD9, CD81), heat shock proteins (HSP90AA1, HSP90AB1), and ALIX. They also included 76% of the top 25 exosome markers registered in Exocarta, which contains molecular data on published and unpublished exosomal studies (http://www.exocarta.org). This result highlights that the evaluated samples included EVs.

**Figure 6.**
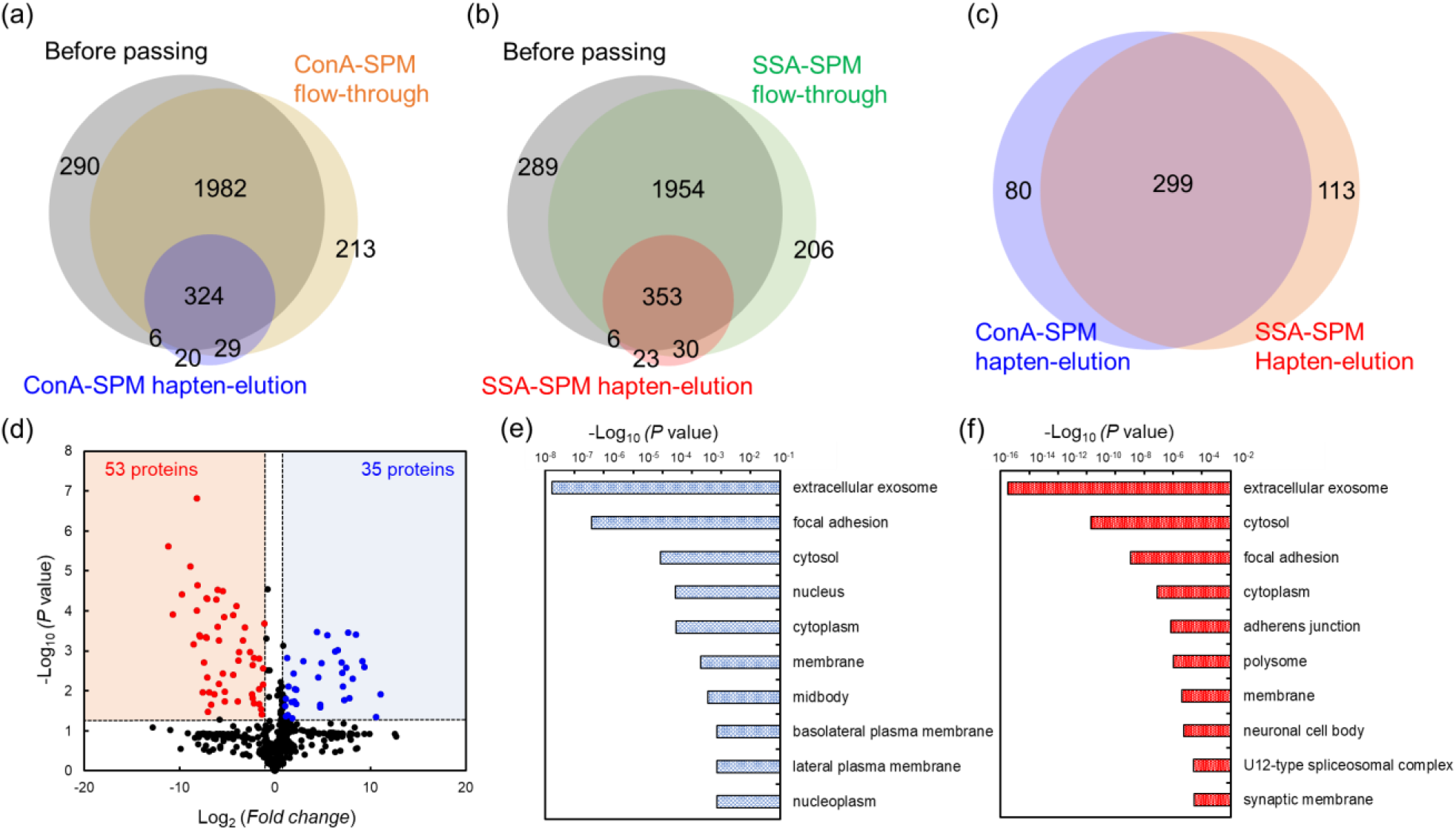
(a), (b), (c) Overlap of proteins quantified in each fraction. (d) Volcano plot showing the - log10 *P*-values as a function of the log_2_ ratios between the EVs from the ConA-SPM and SSA-SPM hapten elutions. Horizontal and vertical axes represent the log_2_ ratio of the fold change and -log10 (Welch’s *t*-test, *P* value), respectively. Blue (red) indicates a more than two-fold increase (decrease) in protein expression in EVs from the SSA-SPM hapten elutions compared to that from the ConA-SPM hapten elutions. Top ten items in cell components from GO enrichment analysis of upregulated proteins in the (e) ConA-SPM and (f) SSA-SPM hapten-elutions. Proteome analyses are performed in triplicate using independent experiments with technical repeats for each sample.

We then calculated Spearman’s rank correlation between EV samples with the intensity based absolute quantification (iBAQ) value, a measure of protein abundance in the exact replicate (Fig. S6, Supporting Information). The correlation coefficients showed that each protein expression level was best correlated with itself across the three trials. The protein abundance clearly differed between the original EVs and EVs in hapten-elution fractions. For example, EVs in the hapten-elution increased the abundance of CD9 relative to CD81 compared to the original EVs. Glycosylation of CD9 is registered in the UniProt database, while CD81 has yet to be reported (Fig. S7 (a), Supporting Information).^[54]^ In addition, EVs eluted from SSA-SPM significantly increased the expression of integrin αv compared to integrin α1, which are well-known membrane proteins similar to tetraspanins (Fig. S7 (b), Supporting Information). These results suggest that EVs can be separated based on their surface glycans.

To investigate the upregulated proteins in the hapten-elution fractions, the fold change values (FC) and *p*-values were calculated using the label-free quantification (LFQ) intensities by Welch’s *t*-test (Fig. S8, Supporting Information). An FC >1 and *p*-value <0.05 were set as the criteria to select upregulated proteins. ConA-SPM and SSA-SPM contained 35 proteins and 53 proteins upregulated in the hapten-elution fractions, respectively. They were depicted in volcano plots. According to Gene Ontology (GO) annotation obtained from the UniProt database using David 6.8 database,^[55]^ the upregulated proteins were assigned different cellular components, and extracellular exosome was annotated as the most over-represented GO terms on both lectin-immobilized SPMs. Furthermore, specific constitutive proteins were also identified in the hapten-elution fractions on each lectin-immobilized SPM (Fig. 6c). The volcano plot showed that the expression of 74 proteins significantly increased on ConA-SPM, whereas the expression of 81 proteins significantly increased on SSA-SPM (Fig. 6d). GO annotation showed these proteins also annotated with extracellular exosome as the most over-represented GO terms (Figs. 6e,f). These results clearly demonstrate that the partial EV subpopulations are enriched by their affinity for the respective lectins and suggests EV heterogeneity based on their surface glycans.

## 3. Conclusion

A novel lectin-affinity separation platform is proposed to classify EVs based on their surface glycan structure. Lectin-immobilized SPMs are effective in the affinity separation for glycoproteins. The separation behavior of mannose-labeled liposomes suggests that SPMs provide a superior recovery rate of liposomal nanoparticles compared to particulate separation media. A simple blocking treatment with PVP for hydrophilization of the surface minimizes the nonspecific adsorption of EVs on SPM without impairing the affinity of the lectins. Finally, our SPMs can enrich EVs with different protein compositions, depending on their affinity with the lectins. Our results provide evidence that EVs have heterogeneity based on their surface glycan structures, even if they are derived from the same cell type and identical size distributions. Our SPMs have potential to become the standard for separating various bio-related nano- and micro-particles. Hence, SPMs should greatly contribute to studies on biological phenomena regulated by EV heterogeneity.

## 4. Experimental Section

### 4.1. Preparation of lectins and protein A immobilized SPMs

An SPM was synthesized as reported previously.^[44]^ The detailed preparation and packing procedures are summarized in Supporting Information. A phosphate buffered salts (PBS) solution was prepared with a PBS tablet (Sigma Aldrich Japan, Tokyo, Japan) into pure water of 100 mL (9.57 mM, pH 7.5). Con-A and SSA (5 mg) were dissolved in the PBS solution of 10 mL. For conditioning the column, acetonitrile (ACN), and deionized water were passed through the SPM at room temperature for 5 mL in each solvent. Each lectin solution (1 mg mL^-1^) was filled into the SPM completely, then the column was incubated at 37 °C for 16 h. The completed column was washed with deionized water for 1 h at 1 mL min^-1^ (ConA-SPM and SSA-SPM). For evaluating the nonspecific adsorption of EVs, the protein A-immobilized SPM was also prepared. The tablet-type SPM was completely soaked and incubated in a protein A solution with PBS (2 mg mL^-1^) at 37 °C for 16 h and shoved into the SPE cartridge (8.0 mm I.D.). Lectins and protein A were purchased from Fujifilm Wako (Osaka, Japan).

### 4.2. Lectin affinity chromatography

We carried out LC analyses with a photodiode array and/or fluorescence detectors by an LC-30 Prominence (Shimadzu Co., Kyoto, Japan). A stepwise gradient mode was employed for LAC evaluations at 25 °C under 0.5 mL min^-1^. Mobile phase A was composed of 2-[4-(2-hydroxyethyl)-1-piperazinyl]-ethanesulfonic acid (HEPES) buffer (20 mM, pH7.4) with 150 mM NaCl, 1 mM MgCl_2_, 1 mM CaCl_2_, and 1 mM MnCl_2_. Mobile phase B was composed of mobile phase A with 200 mM hapten sugar, which can reverse the lectin-carbohydrate interactions and facilitate elution. Methyl *α*-D-mannopyranoside (Tokyo Chemical Industry, Tokyo, Japan) and lactose (Sigma Aldrich Japan) were utilized as hapten sugar to ConA and SSA, respectively. The gradient applied was 100% A for 10 min, 100% B for 10 min, and 100% A for 10 min.

### 4.3. Isolation of EVs from HEK293sus by ultracentrifugation

The human embryonic kidney cell line HEK293sus [CRL-1573; American Type Culture Collection (ATCC)] was cultured in 293 SFM II medium (Thermo Fisher Scientific, MI, USA) supplemented with 2% GlutaMAX™-I (Thermo Fisher Scientific). Cells were grown until ≈70–80% confluence and then cultured in an EV-depleted medium for 48 h. EVs were collected from the resulting culture supernatant by ultracentrifugation to reduce heterogeneity caused by their size. The supernatant was centrifuged at 300 × *g* for 10 min, 2000 × *g* for 10 min, and 10000 × *g* for 30 min at 4 °C. After filtration with 0.22 μm membrane filter, and then ultracentrifuged at 120000 *g* for 100 min at 4 °C. The EV pellets were washed with PBS by re-centrifugation under the same conditions. Protein concentrations were determined using BCA assay (Thermo Fisher Scientific). Protein concentrations of EVs were adjusted to 300 μg mL^-1^.

### 4.4. Evaluation of recovery rates and physical properties of EVs

The detection of proteins and EVs were carried out at 280 nm by the photodiode array. In the SPE evaluation, EVs were labeled with a Cy3 Mono-Reactive dye pack (GE Healthcare Ltd., Tokyo, Japan) for detection by fluorescence microplate readers (Molecular Devices Japan Ltd., Tokyo, Japan). The size distribution of EVs was determined by nano tracking analysis using a NanoSight LM10 (NanoSight, Amesbury, UK) with a blue laser. For analysis, the exosome solution was diluted to about 10^8^–10^9^ particles/mL.

### 4.5. Protein digestion and purification of peptide samples for LC/MS/MS analysis

EVs were dried and digested using the phase-transfer surfactant (PTS)-aided trypsin digestion protocol as described previously.^[56]^ Proteins were reduced with 10 mM dithiothreitol (DTT) (FUJIFILM Wako Pure Chemical Corporation) for 30 min at 37°C, followed by alkylation with 50 mM 2-iodoacetamide (IAA) (FUJIFILM Wako Pure Chemical Corporation) for 30 min at room temperature in the dark. The samples were diluted to 2 M urea with 50 mM ammonium bicarbonate. The proteins were digested with 1 μg lysyl endopeptidase (LysC) (FUJIFILM Wako Pure Chemical Corporation) and 1 μg trypsin (Promega, Tokyo, Japan) overnight at 37°C on a shaking incubator. The resulting peptides were acidified with 0.5% trifluoroacetic acid (TFA, final concentration), and fractionated with a StageTip containing SDB-XC (upper) and SCX (bottom) Empore disk membranes (GL Sciences, Tokyo, Japan).^[57]^ Peptides were washed by 0.1% TFA and 5% ACN and 0.1% TFA and 80% ACN. Then, they eluted from the tip by 500 mM ammonium acetate, 30% ACN and 4% TFA, and 500 mM ammonium acetate and 30% ACN. The sample solution was evaporated in a SpeedVac (Thermo Fisher Scientific) and the residue was resuspended in 0.5% TFA and 4% ACN. Finally, the peptides were desalted again by StageTip with SDB-XC Empore disk membranes and suspended in the loading buffer (0.5% TFA and 4% ACN) for subsequent LC/MS/MS analyses. After digestion, peptide concentration was measured on NanoDrop (Thermo Fisher Scientific) using absorbance at 205 nm and an extinction coefficient of 31.^[58]^

### 4.6. LC/MS/MS analysis

NanoLC/MS/MS analyses were performed on a Q-Exactive (Thermo Fisher Scientific), which was connected to an UltiMate 3000 pump (Thermo Fisher Scientific) and an HTC-PAL autosampler (CTC Analytics). Peptides were separated on pulled in house needle columns (150-mm length, 100 μm inner diameter, 6-μm needle opening) packed with ReproSil-Pur 120 C18-AQ 3-μm RP material (Dr Maisch, Ammerbuch, Germany).^[46]^ The samples were applied by 5-μL full loop injection, and the flow rate was 500 nL/min. Separation was achieved by using a four-step linear gradient of 4 to 10% ACN in 5 min, 10 to 40% MeCN in 60 min, 40 to 99% MeCN in 10 min, and 99% MeCN for 10 min with 0.5%TFA. The electrospray voltage was set to 2.4 kV in the positive mode. The full MS scan was acquired with the mass range of 350-1500 *m/z*, resolution of 70,000, automatic gain control (AGC) target of 3e^6^, and maximum injection time of 100 ms. MS/MS scan was performed by the Top10 method with the resolution of 17,500, the AGC target of 1e^5^, a maximum injection time of 100 ms, and an isolation window of 2.0 Th. The precursor ions were fragmented by higher-energy collisional dissociation with a normalized collision energy of 27%

### 4.7. Database searching

For all experiments, the raw MS data files were analyzed by MaxQuant v2.0.3.0.^[59]^ Peptides and proteins were identified using automated database searching using Andromeda against the human SwissProt Database (version 2022-02, 20,588 protein entries) with a precursor mass tolerance of 20 ppm for the first search and 4.5 ppm for main search and a fragment ion mass tolerance of 20 ppm. The enzyme was set as Trypsin/P with two missed cleavages allowed. Cysteine carbamidomethylation was set as a fixed modification. Methionine oxidation and acetylation on the protein *N*-terminus were set as variable modifications. The search results were filtered with FDR <1% at the peptide spectrum match (PSM) and protein levels. Match-between-run algorithm (MBR) was utilized through the “Identification” subtab in the “Global Parameters” tab of MaxQuant to mitigate the missing value problem. The default settings for MBR were used (0.7 min match window and 20 min alignment time). Proteins that have “Only identified by site”, “potential contaminants” and “reverse sequences” were removed for data analysis. For the missing or zero values, we replaced them with minimum value of that attribute to evaluate relative abundance.

## Acknowledgements

This work was supported by Grants-in-Aid for Scientific Research (KAKENHI; Grants JP21K14652, and JP20K20567), JST Strategic Basic Research Program (CREST; Grant JPMJCR17H2), JST Strategic Basic Research Program (PREST; Grant JPMJPR18H2).

## Author Contributions

Experiments were designed by E. Kanao, T. Kubo, K. Akiyoshi, J. Adachi, K. Otsuka, and Y. Ishihama. SPM columns and mannose-labeled liposomes were prepared by S. Wada and T. Tanigawa. Chromatographic analysis and NTA analysis were operated by S. Wada. EV samples were collected by K. Umezaki. Proteome analyses were operated by E. Kanao, H. Nishida and K. Imami. The manuscript was written by E. Kanao and T. Kubo.

## Data Availability Statement

The MS raw data and analysis files have been deposited at the ProteomeXchange Consortium (http://proteomecentral.proteomexchange.org) via the jPOST partner repository (https://jpostdb.org) with the dataset identifier JPST001529.^[60]^

## Entry for the Table of Contents

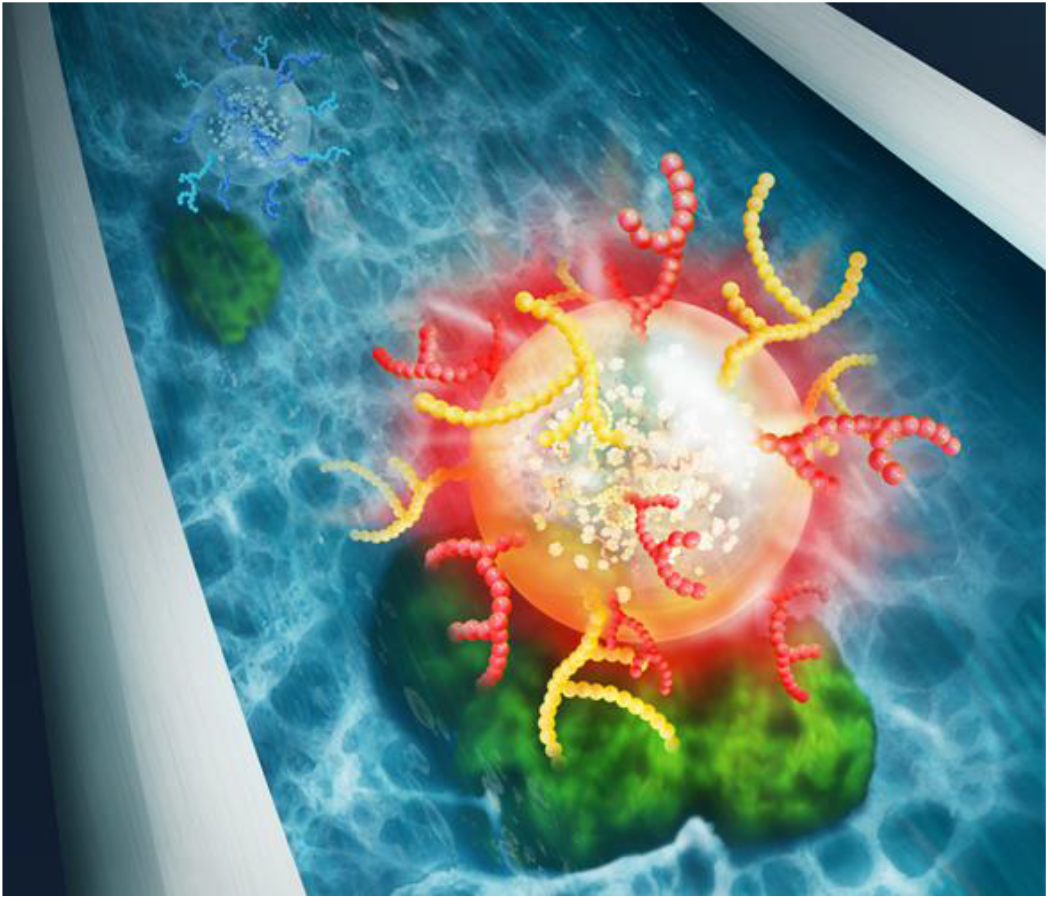

Extracellular vesicles (EVs) are lipid bilayer vesicles that enclose various biomolecules. The potential of a novel lectin-based affinity chromatography (LAC) method to classify EVs based on their glycan structures is demonstrated. Finally, lectin-immobilized SPMs are employed to classify EVs based on the surface glycan structures and demonstrate different subpopulations by proteome profiling.

## References

[1] R. Kalluri, S. LeBleu Valerie, Science 2020, 367, eaau6977.

[2] G. Raposo, W. Stoorvogel, J. Cell Biol. 2013, 200, 373.

[3] H. Valadi, K. Ekström, A. Bossios, M. Sjöstrand, J. J. Lee, J. O. Lötvall, Nat. Cell Biol. 2007, 9, 654.

[4] O. G. de Jong, M. C. Verhaar, Y. Chen, P. Vader, H. Gremmels, G. Posthuma, R. M. Schiffelers, M. Gucek, B. W. M. van Balkom, J. Extracell. Vesicles 2012, 1, 18396.

[5] V. S. LeBleu, R. Kalluri, Trends Cancer 2020, 6, 767.

[6] M. Nawaz, G. Camussi, H. Valadi, I. Nazarenko, K. Ekström, X. Wang, S. Principe, N. Shah, N. M. Ashraf, F. Fatima, L. Neder, T. Kislinger, Nat. Rev. Urol. 2014, 11, 688.

[7] M. Niu, Y. Li, G. Li, L. Zhou, N. Luo, M. Yao, W. Kang, J. Liu, Eur. J. Neurol. 2020, 27, 967.

[8] N. Osier, V. Motamedi, K. Edwards, A. Puccio, R. Diaz-Arrastia, K. Kenney, J. Gill, Mol. Neurobiol. 2018, 55, 9280.

[9] J. Dai, Y. Su, S. Zhong, L. Cong, B. Liu, J. Yang, Y. Tao, Z. He, C. Chen, Y. Jiang, Signal Transduct. Target. Ther. 2020, 5, 145.

[10] W. Guo, Y. Li, W. Pang, H. Shen, Mol. Ther. 2020, 28, 1953.

[11] M. N. Huda, M. Nafiujjaman, I. G. Deaguero, J. Okonkwo, M. L. Hill, T. Kim, M. Nurunnabi, ACS Biomater. Sci. Eng. 2021, 7, 2106.

[12] D. K. Jeppesen, A. M. Fenix, J. L. Franklin, J. N. Higginbotham, Q. Zhang, L. J. Zimmerman, D. C. Liebler, J. Ping, Q. Liu, R. Evans, W. H. Fissell, J. G. Patton, L. H. Rome, D. T. Burnette, R. J. Coffey, Cell 2019, 177, 428.

[13] Y. Meng, W. Lu, E. Guo, J. Liu, B. Yang, P. Wu, S. Lin, T. Peng, Y. Fu, F. Li, Z. Wang, Y. Li, R. Xiao, C. Liu, Y. Huang, F. Lu, X. Wu, L. You, D. Ma, C. Sun, P. Wu, G. Chen, J. Hematol. Oncol. 2020, 13, 75.

[14] A. Bobrie, M. Colombo, S. Krumeich, G. Raposo, C. Théry, J. Extracell. Vesicles 2012, 1, 18397.

[15] J. Kowal, G. Arras, M. Colombo, M. Jouve, J. P. Morath, B. Primdal-Bengtson, F. Dingli, D. Loew, M. Tkach, C. Théry, Proc. Natl. Acad. Sci. USA 2016, 113, E968.

[16] F. G. Kugeratski, K. Hodge, S. Lilla, K. M. McAndrews, X. Zhou, R. F. Hwang, S. Zanivan, R. Kalluri, Nat. Cell Biol. 2021, 23, 631.

[17] S. Matsumura, T. Minamisawa, K. Suga, H. Kishita, T. Akagi, T. Ichiki, Y. Ichikawa, K. Shiba, J. Extracell. Vesicles 2019, 8, 1579541.

[18] W. Nakai, T. Yoshida, D. Diez, Y. Miyatake, T. Nishibu, N. Imawaka, K. Naruse, Y. Sadamura, R. Hanayama, Sci. Rep. 2016, 6, 33935.

[19] G. M. Barton, J. C. Kagan, Nat. Rev. Immunol. 2009, 9, 535.

[20] A. M. Bourgonje, A. C. Navis, J. T. G. Schepens, K. Verrijp, L. Hovestad, R. Hilhorst, S. Harroch, P. Wesseling, W. P. J. Leenders, W. J. A. J. Hendriks, Oncotarget 2014, 5, 8690.

[21] S. Gurung, D. Perocheau, L. Touramanidou, J. Baruteau, Cell Commun. Signal. 2021, 19, 47.

[22] K. O’Brien, K. Breyne, S. Ughetto, L. C. Laurent, X. O. Breakefield, Nat. Rev. Mol. Cell Biol. 2020, 21, 585.

[23] J. Ratelade, A. S. Verkman, Int. J. Biochem. Cell Biol. 2012, 44, 1519.

[24] B. H. Sung, A. von Lersner, J. Guerrero, E. S. Krystofiak, D. Inman, R. Pelletier, A. Zijlstra, S. M. Ponik, A. M. Weaver, Nat. Commun. 2020, 11, 2092.

[25] K. J. Svensson, H. C. Christianson, A. Wittrup, E. Bourseau-Guilmain, E. Lindqvist, L. M. Svensson, M. Mörgelin, M. Belting, J. Biol. Chem. 2013, 288, 17713.

[26] K. Ohtsubo, J. D. Marth, Cell 2006, 126, 855.

[27] C. Reily, T. J. Stewart, M. B. Renfrow, J. Novak, Nat. Rev. Nephrol. 2019, 15, 346.

[28] J. Costa, M. Gatermann, M. Nimtz, S. Kandzia, M. Glatzel, H. S. Conradt, Anal. Chem. 2018, 90, 7871.

[29] J. Gomes, P. Gomes-Alves, S. B. Carvalho, C. Peixoto, P. M. Alves, P. Altevogt, J. Costa, Biomolecules 2015, 5.

[30] N. Nishida-Aoki, N. Tominaga, N. Kosaka, T. Ochiya, J. Extracell. Vesicles 2020, 9, 1713527.

[31] M. Surman, D. Hoja-Łukowicz, S. Szwed, A. Drożdż, E. Stępień, M. Przybyło, Life Sci. 2018, 207, 395.

[32] C. Williams, R. Pazos, F. Royo, E. González, M. Roura-Ferrer, A. Martinez, J. Gamiz, N.-C. Reichardt, J. M. Falcón-Pérez, Sci. Rep. 2019, 9, 11920.

[33] J. Q. Gerlach, A. Krüger, S. Gallogly, S. A. Hanley, M. C. Hogan, C. J. Ward, L. Joshi, M. D. Griffin, PLoS One 2013, 8, e74801.

[34] A. Shimoda, S.-i. Sawada, Y. Sasaki, K. Akiyoshi, Sci. Rep. 2019, 9, 11497.

[35] A. Shimoda, R. Miura, H. Tateno, N. Seo, H. Shiku, S.-i. Sawada, Y. Sasaki, K. Akiyoshi, Small Methods 2021, n/a, 2100785.

[36] A. Shimoda, Y. Tahara, S.-i. Sawada, Y. Sasaki, K. Akiyoshi, Biochem. Biophys. Res. Commun. 2017, 491, 701.

[37] I. Fakla, A. Hever, J. Molnár, J. Fischer, Anticancer Res. 1998, 18, 3107.

[38] Y. Liu, D. Fu, Y. Xiao, Z. Guo, L. Yu, X. Liang, Anal. Method 2015, 7, 25.

[39] B. F. Mann, A. K. P. Mann, S. E. Skrabalak, M. V. Novotny, Anal. Chem. 2013, 85, 1905.

[40] A. Monzo, G. K. Bonn, A. Guttman, Trends Anal. Chem. 2007, 26, 423.

[41] S. Jung, S. Ehlert, M. Pattky, U. Tallarek, J. Chromatogr. A 2010, 1217, 696.

[42] C. A. Rimmer, C. R. Simmons, J. G. Dorsey, J. Chromatogr. A 2002, 965, 219.

[43] S. Tajima, S. Yokoyama, A. Yamamoto, J. Biol. Chem. 1983, 258, 10073.

[44] K. Kubota, T. Kubo, T. Tanigawa, T. Naito, K. Otsuka, Sci. Rep. 2017, 7, 178.

[45] S. Warwood, A. Byron, M. J. Humphries, D. Knight, J. Proteomics 2013, 85, 160.

[46] Y. Ishihama, J. Rappsilber, J. S. Andersen, M. Mann, J. Chromatogr. A 2002, 979, 233.

[47] N. Itoh, A. Kimoto, E. Yamamoto, T. Higashi, T. Santa, T. Funatsu, M. Kato, J. Chromatogr. A 2017, 1484, 34.

[48] P. Lundahl, C.-M. Zeng, C. Lagerquist Hägglund, I. Gottschalk, E. Greijer, J. Chromatogr. B 1999, 722, 103.

[49] L. Hagel, M. Östberg, T. Andersson, J. Chromatogr. A 1996, 743, 33.

[50] F. A. W. Coumans, A. R. Brisson, E. I. Buzas, F. Dignat-George, E. E. E. Drees, S. El-Andaloussi, C. Emanueli, A. Gasecka, A. Hendrix, A. F. Hill, R. Lacroix, Y. Lee, T. G. van Leeuwen, N. Mackman, I. Mäger, J. P. Nolan, E. van der Pol, D. M. Pegtel, S. Sahoo, P. R. M. Siljander, G. Sturk, O. de Wever, R. Nieuwland, Circ. Res. 2017, 120, 1632.

[51] E. G. Evtushenko, D. V. Bagrov, V. N. Lazarev, M. A. Livshits, E. Khomyakova, PLoS One 2021, 15, e0243738.

[52] K. W. Witwer, E. I. Buzás, L. T. Bemis, A. Bora, C. Lässer, J. Lötvall, E. N. Nolte-’t Hoen, M. G. Piper, S. Sivaraman, J. Skog, C. Théry, M. H. Wauben, F. Hochberg, J. Extracell. Vesicles 2013, 2, 20360.

[53] K. Ishihara, H. Nomura, T. Mihara, K. Kurita, Y. Iwasaki, N. Nakabayashi, J. Biomed. Mater. Res. 1998, 39, 323.

[54] S. Charrin, S. Manié, M. Oualid, M. Billard, C. Boucheix, E. Rubinstein, FEBS Lett. 2002, 516, 139.

[55] D. W. Huang, B. T. Sherman, R. A. Lempicki, Nat. Protoc. 2009, 4, 44.

[56] T. Masuda, M. Tomita, Y. Ishihama, J. Proteome Res. 2008, 7, 731.

[57] J. Adachi, K. Hashiguchi, M. Nagano, M. Sato, A. Sato, K. Fukamizu, Y. Ishihama, T. Tomonaga, Anal. Chem. 2016, 88, 7899.

[58] R. K. Scopes, Anal. Biochem. 1974, 59, 277.

[59] J. Cox, M. Mann, Nat. Biotechnol. 2008, 26, 1367.

[60] S. Okuda, Y. Watanabe, Y. Moriya, S. Kawano, T. Yamamoto, M. Matsumoto, T. Takami, D. Kobayashi, N. Araki, A. C. Yoshizawa, T. Tabata, N. Sugiyama, S. Goto, Y. Ishihama, Nucleic Acids Res. 2017, 45, D1107.

